# SimRVSequences: an R package to simulate genetic sequence data for pedigrees

**DOI:** 10.1101/534552

**Authors:** Christina Nieuwoudt, Angela Brooks-Wilson, Jinko Graham

## Abstract

1

**Summary:** Family-based studies have several advantages over case-control studies for finding causal rare variants for a disease; these include increased power, smaller sample size requirements, and improved detection of sequencing errors. However, collecting suitable families and compiling their data is time-consuming and expensive. To evaluate methodology to identify causal rare variants in family-based studies, one can use simulated data. For this purpose we present the `R` package `SimRVSequences`. Users supply a sample of pedigrees and single-nucleotide variant data from a sample of unrelated individuals representing the pedigree founders. Users may also model genetic heterogeneity among families. For ease of use, `SimRVSequences` offers methods to import and format single-nucleotide variant data and pedigrees from existing software.

**Availability and Implementation:** `SimRVSequences` is available as a library for `R` *≥* 3.5.0 on the comprehensive R archive network.

## 2 Introduction

Family-based studies are comprised of pedigrees, which may be small or may include many individuals and span several generations. Families may belong to a population cohort, or they may be ascertained on the basis of disease-affected relatives. When the disease of interest is rare, the possibility of observing pedigrees with multiple disease-affected relatives by chance alone is small. However, for diseases with a sporadic component, study samples may include both genetic and sporadic pedigrees, and pedigrees with both genetic and sporadic cases. To evaluate methodology to identify causal rare variants (*c*RVs) in family-based studies of disease, we have developed the `R` package `SimRVSequences`.

Family-based studies can be more attractive than case-control studies because they have more power to detect causal rare variants (Wijsman, 2012). Even so, family-based studies may fail to identify causal rare variants owing to genetic heterogeneity, which occurs when different loci predispose to the same disease (Nussbaum et al., 2007). The disease-predisposing variants may be restricted to a single gene or they may be contained in a pathway, or a set of related genes. `SimRVSequences` allows users to model genetic heterogeneity and has added functionality to simulate genetic heterogeneity in a pathway.

Several approaches have been proposed to simulate genetic sequence data for pedigrees; e.g. `sim1000G` (Dimitromanolakis et al., 2019) and RarePedSim (Li et al., 2015). Both of these programs reduce user burden by simulating founder haplotypes for pedigree data; however, neither can accommodate large simulations. `SimRVSequences`, on the other hand, can accommodate genome-wide, exon-only simulations. To our knowledge, `SimRVSequences` is the only `R` package that simulates genetic sequence data conditionally given the *c*RV status of family members. This feature allows users to specify causal rare variants prior to simulating genetic data in pedigrees. For additional information regarding the features of these programs please refer to the supplementary table.

## 3 Description

`SimRVSequences` is a publicly-available, platform-independent R package for `R` *≥* 3.5.0, which can be obtained from the comprehensive R archive network (CRAN) (R Core Team, 2013). To simulate sequence data for a family-based study with `SimRVSequences`, users must provide:

- a sample of ascertained pedigrees, and
- single-nucleotide variant (SNV) data from unrelated individuals representing the population of pedigree founders.

We now describe each of the user-supplied items in detail followed by a brief description of the model to simulate genetic data for pedigrees.

### 3.1 Pedigree Data

`SimRVSequences` can accept pedigree data from any source provided that pedigrees are properly formatted. In particular, users who wish to simulate the segregation of known *c*RVs must specify the *c*RV status of each pedigree member. For ease of use, we note that pedigrees simulated by the `R` package `SimRVPedigree` are appropriately formatted for `SimRVSequences`. `SimRVPedigree` simulates pedigrees dynamically by way of a competing risk model and makes use of agespecific hazard rates of disease-onset and death provided by the user. At the individual level, disease onset is influenced by the presence (or absence of) a *c*RV, by way of a proportional hazards model. At the family level, `SimRVPedigree` models complex ascertainment so that simulated pedigrees are representative of sampled pedigrees (Nieuwoudt et al., 2018).

Properly formatted pedigrees are data frames that include the following variables.

1. `FamID`: family identification number
2. `ID`: individual identification number
3. `sex`: sex identification variable: `sex = 0` for males, and `sex = 1` females.
4. `dadID`: identification number of father
5. `momID`: identification number of mother
6. `affected`: disease-affection status: `affected = TRUE` if individual has developed disease, and `FALSE` otherwise.
7. `DA1`: paternally inherited allele at the *c*RV locus: `DA1 = 1` if the *c*RV is inherited, and `0` otherwise.
8. `DA2`: maternally inherited allele at the *c*RV locus: `DA2 = 1` if the *c*RV is inherited, and `0` otherwise.

Please note, if the variables `DA1` and `DA2` are not provided, the pedigree is assumed to be fully-sporadic, i.e. `DA1 = DA2 = 0` for all members. For additional information regarding the required pedigree format please refer to the supplementary file.

### 3.2 Single-Nucleotide Variant Data

By convention, each individual in a pedigree can be classified as a founder or a non-founder. Founders are individuals whose parents are missing, while non-founders have both a mother and a father represented in the pedigree. Users are required to provide the haplotypes for the founders in the form of SNV data.

`SimRVSequences` can accept SNV data obtained from various sources (e.g. 1000 genomes project); provided they are properly formatted. For ease of use, `SimRVSequences` includes methods to import and format SNV data simulated by the freely-available software SLiM (Haller and Messer, 2017). We have found SLiM to be the most practical for simulation of genome-wide, exon-only SNV data in a large sample of unrelated individuals.

Users interested in simulating exon-only sequence data with SLiM will enjoy the additional functionality provided by `SimRVSequences`. With SLiM, users may specify a recombination map to simulate mutations over non-contiguous genomic regions such as the exons (e.g. Harris and Nielsen (2016)). `SimRVSequences` may be used to create the recombination map to simulate exon-only data in SLiM. To create the recombination map, users provide a map of exons and specify the genome-wide, per-site, per-generation recombination and mutation rates. If desired, a map of exons from the UCSC Genome Browser (Kent et al., 2002) is available for this task. By default, we set the recombination rate to be 1 *×* 10^-8^ per-site per-generation (Harris and Nielsen, 2016), and assume that all mutations in exons are neutral and arise at a rate of 1 *×* 10^-8^ per-site per-generation (Harris and Nielsen, 2016; 1000 Genomes Project, 2010). Users may alter the default settings, however, to create a variety of recombination maps for SLiM.

After simulating the SNV data, `SimRVSequences` may be used to import SLiM data into `R`. In addition to importing the SLiM data to `R`, `SimRVSequences` will format the data appropriately. For additional information regarding the required format for SNV data please refer to the supplementary file.

### 3.3 Pathway Implementation and Selection of Founder Haplotypes

Upon importing and formatting SNV data, the user must specify which mutations will be modelled as *c*RVs. We allow users to model genetic heterogeneity through the specification of a pool of *c*RVs from a pathway. Users may also implement allelic heterogeneity among families by selecting *c*RVs in the same gene. Each pedigree is assumed to segregate at most one *c*RV. Thus, for each pedigree segregating a causal variant, we sample a single *c*RV from the user-specified pool of *c*RVs, according to their relative frequency, with replacement. Hence, different families may segregate different *c*RVs.

After identifying the familial rare variant, the haplotypes for each pedigree founder are sampled from the user-supplied distribution of haplotypes conditioned on the founder’s *c*RV status at the familial disease locus. Thus, we ensure that the *c*RV is introduced by the appropriate founder.

For additional information regarding the specification of *c*RVs please refer to the supplementary material.

### 3.4 Simulation of Sequence Data for Non-founders

To simulate sequence data for non-founders we perform a conditional gene-drop, which can be described as a two-part process. The first step involves simulating the formation of gametes from parental haplotypes, and the second involves conditional inheritance of a gamete given the offspring’s *c*RV status at the familial disease locus.

To simulate formation of gametes we use the model proposed by Voorrips and Maliepaard (2012) for recombination with chiasmata interference. Although the proposed model can be applied to tetraploids, in `SimRVSequences` we restrict attention to diploid organisms.

Given a parent-offspring pair, we simulate the formation of four gamete cells from the genetic material of the parent. To determine which gamete is transmitted from parent to offspring, we employ a conditional gene-drop algorithm which depends upon the *c*RV status of the parent and the offspring. This algorithm is described as follows:

- Case 1: If both the parent and offspring carry the *c*RV, we sample the offspring’s gamete from the parental gametes that carry the *c*RV with equal probability. For example, in the father to offspring transmission in Figure 1, we choose from the paternal gametes represented by the top two chromatids.

**Figure 1:**
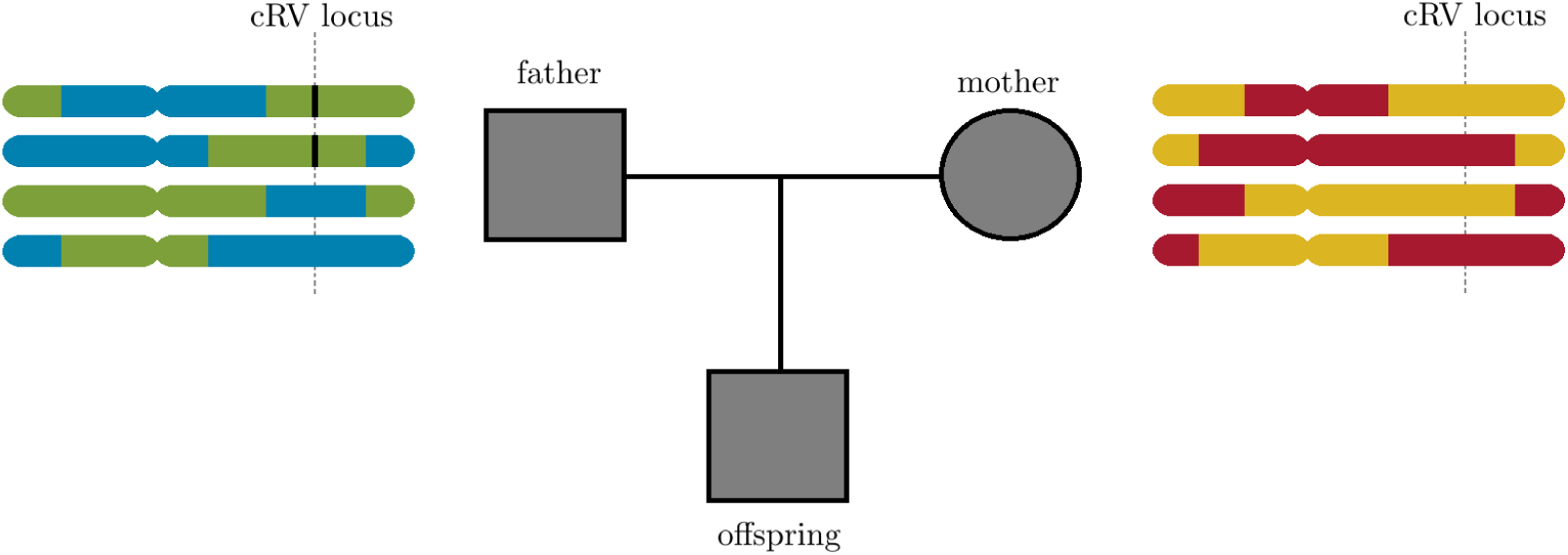
A family trio with possible maternal and paternal recombinant chromatids for a diploid organism with a single set of homologous chromosomes. The *c*RV locus is indicated by a dashed vertical line over the chromatids, and the *c*RV is indicated by a black vertical bar within the chromatids. The top two chromatids of the father contain the only copies of the *c*RV.
- Case 2: If the parent carries the *c*RV but the offspring does not, we sample the offspring’s gamete from the parental gametes that do not carry the *c*RV with equal probability. For example, in the father to offspring transmission in Figure 1, we choose from the paternal gametes represented by the bottom two chromatids.
- Case 3: If a parent is not a carrier of the *c*RV, we sample the offspring’s gamete from all possible parental gametes with equal probability. For example, this would be the case in the mother to offspring transmission in Figure 1, since she is not a carrier of the *c*RV.

To simulate genetic data in an entire pedigree, we repeat this process, forwards-in-time, for each parent-offspring pair to ensure the correct genetic variability among relatives.

## 4 Method Overview

The methods provided by `SimRVSequences` are described as follows.

- Methods to assist with creating the recombination map for SLiM:
  - `combine exons`: This function will be useful to users who would like to combine overlapping exon observations into a single observation. Upon supplying a data frame which contains overlapping exon segments, this function will return a data frame with combined segments.
  - `create slimMap`: This function is used both before and after simulating SLiM data. Prior to simulating the SNV data with SLiM, this function can be used to create the recombination map required by SLiM to simulate exon-only SNV data (see section 3.1 of the supplement). Additionally, if this function was used to create the recombination map users may then supply the output of this function to `read slim` to remap mutations to their correct locations when importing SLiM data to `R` (see section 3.2 of the supplement).
- Method to assist with importing SLiM data to `R`.
  - `read slim`: This function is used to import to `R` data stored to a .txt file by SLiM’s outputfull() method (see section 3.2 of the supplement).
- Methods to assist with simulation and basic manipulation of SNV data for pedigrees
  - `sim RVstudy`: This function is used to simulate the SNV data for a sample of pedigrees (see section 5.1 of the supplement).
  - `summary.famStudy`: This function is used to obtain a summary of the SNVs shared by the disease-affected relatives in each pedigree (see section 5.3 of the supplement).

## 5 Example

We demonstrate the procedure to simulate sequence data for a sample of pedigrees given properly formatted inputs. After installing `SimRVSequences` we load the package with the `library` function.

~~~
library(SimRVSequences)
~~~

We simulate sequence data using the `sim RVstudy` function. This function has three required inputs:

- `ped files` A data frame of pedigree data for all families in the study.
- `haplos` A sparse matrix of haplotypes for a sample of unrelated individuals.
- `SNV map` A data frame which describes the mutations in `haplos`; that is, each row of `SNV map` catalogues information about the mutation in the corresponding column of `haplos`.

For additional information regarding the optional inputs for the `sim RVstudy` function please refer to the supplementary material.

Assume that the pedigree data frame is stored as `study peds`; the matrix of haplotypes is stored as `hap mat`; and the mutation data is stored as `mut dat`. To simulate the sequence data for the pedigrees in `ped files` we use the `sim RVstudy` command as follows.

~~~
   study seq <-sim RVstudy(ped files = study peds,
                               haplos = hap mat,
                               SNV map = mut dat)
~~~

As displayed in the command above, the output of the `sim RVstudy` function has been stored as `study seq`. To determine the class of `study seq` we use the `class` function.

~~~
   class(study seq)
   [1] “famStudy” “list”
~~~

From the output above, we see that `sim RVstudy` returns a object of class `famStudy` that inherits from the `list` class. Objects of class `famStudy` are lists that contain the following four items:

- a data frame named `ped files`.
- a sparse matrix named `ped haplos`,
- a data frame named `haplo map`,
- and a data frame named `SNV map`.

The item `ped files` is a data frame containing the pedigree data for the individuals with simulated sequence data. The sparse matrix `ped haplos` contains the simulated haplotypes for each individual in `ped files`. The `haplo map` data frame is used to map the haplotypes (i.e. rows) in `ped haplos` to the individuals in `ped files`. Finally, the output `SNV map` catalogs the SNVs, i.e. columns, in `ped haplos`. For a extensive description of objects of class `famStudy` please refer to the supplementary material.

## 6 Discussion

Although gene-dropping can be computationally expensive, `SimRVSequences` stores and extracts haplotypes efficiently (Cheng et al., 2015; Voorrips and Maliepaard, 2012) for improved performance in `R`. Simulating the transmission of 178,430 rare SNVs across the 22 human autosomes for the disease-affected relatives in one pedigree requires, on average, 0.5 seconds on a Windows OS with an i7-4790 @ 3.60GHz and 12 GB of RAM. Due to data-allocation limitations in `R`, users who wish to simulate more than 250,000 SNVs may need to run chromosome-specific analyses.

We emphasize that, while `SimRVSequences` can model genetic heterogeneity among families, we assume that within a family genetic cases are due to a single causal SNV. Hence, `SimRVSequences` is not appropriate for users who wish to simulate the transmission of multiple causal SNVs in the same pedigree. Furthermore, `SimRVSequences` does not allow for simulation of sequence data in sex-chromosomes.

## 7 Conclusion

Simulated sequence data for family-based studies is essential to evaluate methodology to identify causal rare variants. `SimRVSequences` allows for efficient simulation of SNV data for a sample of pedigrees in `R`. Users may model known genetic heterogeneity and incorporate pedigrees that include individuals with sporadic disease in simulation. With this tool in hand, researchers may evaluate a variety of methods to identify *c*RVs in ascertained pedigrees.

