## Supplementary Material for "SimRVSequences: an R package to simulate genetic sequence data for pedigrees"

### SimRVSequences

*2019-05-13*

**6 `SimRVSequences` Model** **18**

**7 Method Overview** **19**

**8 Timing** **20**

**9 References** **20**

**10 Appendix** **21**

### 1 Introduction

Compared to case-control studies, family-based studies offer more power to detect causal rare variants, require smaller sample sizes, and detect sequencing errors more accurately [12]. However, data collection for family-based studies is both time-consuming and expensive. Furthermore, evaluating methodology to identify causal rare variants requires simulated data. In this vignette we illustrate how the methods provided by `SimRVSequences` may be used to to simulate sequence data for family-based studies of diploid organisms.

Please note that `SimRVSequences` does NOT allow users to simulate genotypes conditional on phenotype. Instead, `SimRVSequences` simulates haplotypes conditional on the genotype at a single causal rare variant (cRV), via the following algorithm.

1. For each pedigree, we sample a cRV from a pool of SNVs specified by the user (see section 3.3); this is the familial cRV.
2. After identifying the familial cRV we sample founder haplotypes from haplotype data conditional on the founder's cRV status at the familial cRV.

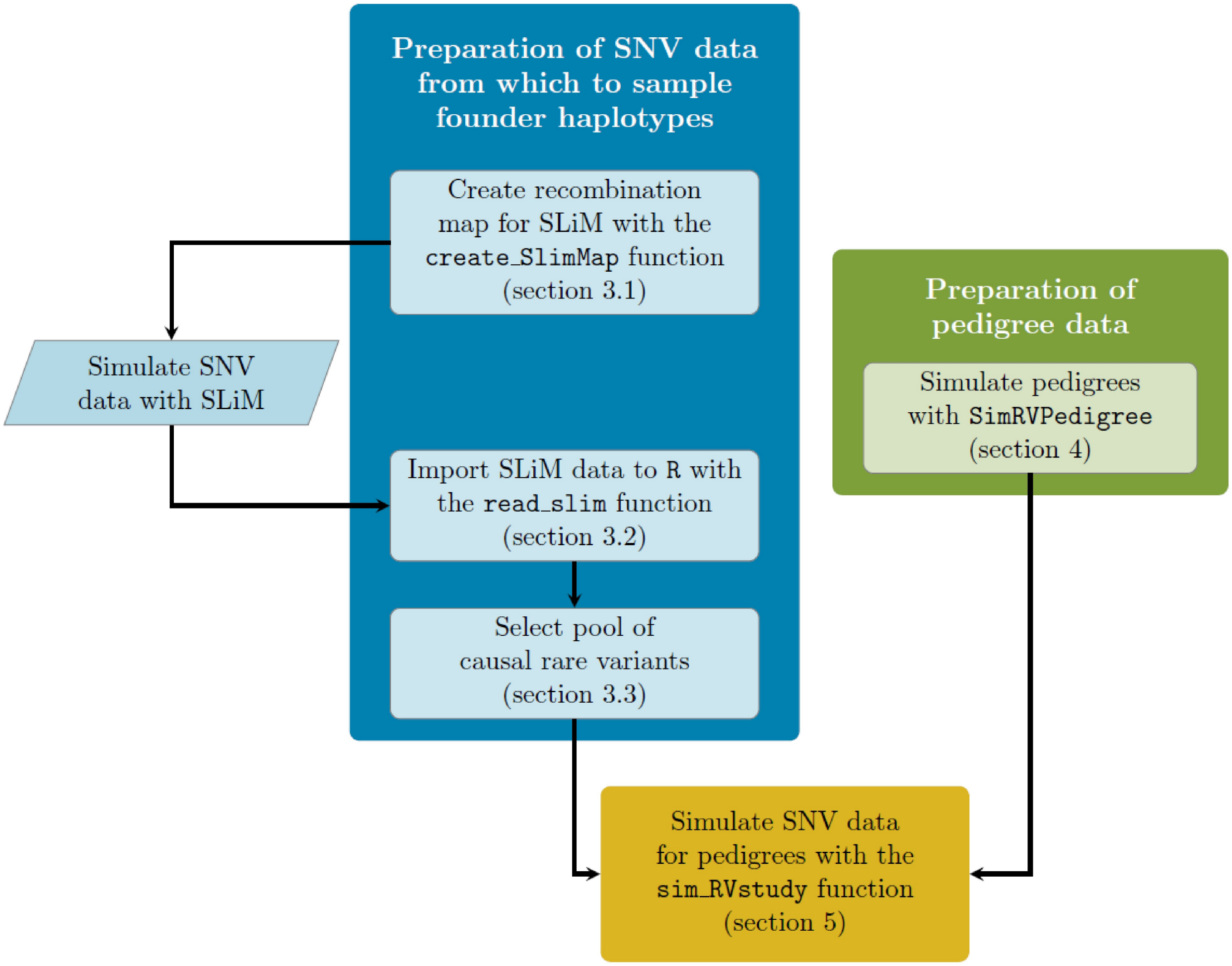

Figure 1: **Figure 1.** Data preparation flow chart. Tasks completed in `R` are enclosed in rectangular frames while the SLiM simulation component is enclosed in a parallelogram.

3. Proceeding forward in time, from founders to more recent generations, for each parent/offspring pair we
    1. simulate recombination and formation of parental gametes (see section 6), and then
    2. perform a conditional gene drop to model inheritance of the cRV (see section 6).

#### 2 User Requirements

To simulate sequence data for a family-based study users are expected to provide:

1. a sample of ascertained pedigrees, and
2. single-nucleotide variant (SNV) data from a sample of unrelated individuals representing the population of pedigree founders.

Presently, we streamline this process for pedigrees simulated by the `R` package `SimRVPedigree` [8] and exon-only SNV data produced by SLiM [3]. The flow chart in Figure 1 illustrates the preparation required for each of the above inputs. We note that simulation of SNV data with SLiM is NOT accomplished in `R`. However, we do provide pre- and post-processing methods to assist with importing and formatting the SNV data simulated by SLiM.

In section 3 we demonstrate how the methods provided by `SimRVSequences` can aid in the preparation of the SNV data, and in section 4 we briefly review simulating pedigree data using the `SimRVPedigree` package.

### 3 Prepare Single Nucleotide Variant Data

Before we can simulate genetic data for a sample of pedigrees, we require single-nucleotide variant (SNV) data for the pedigree founders. Several publicly available programs can aid in this task (e.g. msprime and fastsimcoal), however, we have found the Eidos program SLiM [3] to be the most practical for simulation of genome-wide, exon-only SNV data for a large number of unrelated individuals.

One of SLiM's many features is that it simulates recombination hotspots using a user-specified recombination map. This recombination map may be utilized to simulate SNVs over unlinked regions (i.e. in different chromosomes) or in linked but non-contiguous regions (i.e in exon-only data). In section 3.1 we demonstrate how the `create_slimMap` function may be used to generate the recombination map required by SLiM to simulate exon-only SNV data over the entire human genome.

After simulating SNV data with SLiM, users may import the simulated data to `R` using the `read_slim` function. This task is demonstrated in section 3.2.

Finally, in section 3.3 we demonstrate how users may implement a pathway approach by specifying a pool of rare variants from which to select causal familial variants.

#### 3.1 Create Recombination Map to Simulate Genome-Wide, Exon-Only Data with SLiM

The `SimRVSequences` function `create_slimMap` simplifies the task of creating the recombination map required by SLiM to simulate exon-only SNV data. The `create_slimMap` function has three arguments:

1. `exon_df`: A data frame that contains the positions of each exon to simulate. This data frame must contain the variables `chrom`, `exonStart`, and `exonEnd`. *We expect that `exon_df` does not contain any overlapping segments. Prior to supplying the exon data to `create_slimMap` users must combine overlapping exons into a single observation. Our `combine_exons` function may be used for this task (see appendix for details).*
2. `mutation_rate`: the per-site per-generation mutation rate, assumed to be constant across the genome. By default, `mutation_rate= 1E-8`, as in [4].
3. `recomb_rate`: the per-site per-generation recombination rate, assumed to be constant across the genome. By default, `recomb_rate= 1E-8`, as in [4].

We will illustrate usage of `create_slimMap` with the `hg_exons` data set.

```r
# load the SimRVSequences library
library(SimRVSequences)

# load the hg_exons dataset
data("hg_exons")

# print the first four rows of hg_exons
head(hg_exons, n = 4)
```

```
##   chrom exonStart exonEnd                  NCBIref
## 1     1     11874   12227              NR_046018.2
## 2     1     12613   12721              NR_046018.2
## 3     1     13221   14829 NR_046018.2, NR_024540.1
## 4     1     14970   15038              NR_024540.1
```

As seen in the output above, the `hg_exons` data set catalogs the position of each exon in the 22 human autosomes. The data contained in `hg_exons` were collected from the hg 38 reference genome with the UCSC Genome Browser [5, 6]. The variable `chrom` is the chromosome on which the exon resides, `exonStart` is the position of the first base pair in the exon, and `exonEnd` is the position of the last base pair in the exon. The variable `NCBIref` is the NCBI reference sequence accession numbers for the coding region in which the exon resides. In `hg_exons` overlapping exons have been combined into a single observation. When exons from genes with different NCBI accession numbers have been combined the variable `NCBIref` will contain multiple accession numbers separated by commas. We note that different accession numbers may exist for transcript variants of the same gene.

Since the exons in `hg_exons` have already been combined into non-overlapping segments, we supply this data frame to `create_slimMap`.

```
# create recombination map for exon-only data using the hg_exons dataset
s_map <- create_slimMap(exon_df = hg_exons)

# print first four rows of s_map
head(s_map, n = 4)
```

```
##   chrom segLength  recRate mutRate  exon simDist endPos
## 1     1     11873 0.00e+00   0e+00 FALSE       1      1
## 2     1       354 1.00e-08   1e-08  TRUE     354    355
## 3     1       385 3.85e-06   0e+00 FALSE       1    356
## 4     1       109 1.00e-08   1e-08  TRUE     109    465
```

The `create_slimMap` function returns a data frame with several variables:

1. `chrom`: the chromosome number.
2. `segLength`: the length of the segment in base pairs. *We assume that segments contain the positions listed in `exonStart` and `exonEnd`. Therefore, for a combined exon segment, `segLength` is calculated as `exonEnd - exonStart + 1`.*
3. `recRate`: the per-site per-generation recombination rate. Following [4], segments between exons on the same chromosome are simulated as a single base pair with `rec_rate` equal to recombination rate multiplied by the number of base pairs in the segment. For each chromosome, a single site is created between the last exon on the previous chromosome and the first exon of the current chromosome. This site will have recombination rate 0.5 to accommodate unlinked chromosomes.
4. `mutRate`: the per-site per-generation mutation rate. Since we are interested in exon-only data, the mutation rate outside exons is set to zero.
5. `exon`: logical variable, `TRUE` if segment is an exon and `FALSE` if not an exon.
6. `simDist`: the simulated exon length, in base pairs. When `exon = TRUE`, `simDist = segLength`; however, when `exon = FALSE`, `simDist = 1` since segments between exons on the same chromosome are simulated as a single base pair.
7. `endPos`: The simulated end position, in base pairs, of the segment.

The first row in the output above contains information about the genetic segment **before the first exon** on chromosome 1. Referring back to the `hg_exons` data we see that the first exon begins at position 11,874. Since the region before the first exon contains 11,873 base pairs `segLength = 11873` for this region. To avoid unnecessary computation in SLiM, `simDist = 1` so that this non-exonic region is simulated as a single base pair. The recombination rate for this segment is set to zero. However, the first non-exon segment for the next chromosome will be 0.5 so that chromosomes are unlinked. The mutation rate for this non-exon segment is set to zero since we are interested in exon-only data.

The second row of the output catalogs information for the first exon on chromosome 1. This exon contains 354 base pairs, hence `simDist = 354` for this segment. The per-site per-generation mutation and recombination rates for this exon are both 1E-08.

The third row contains the information for the second non-exon segment on chromosome 1. This segment contains 385 base pairs. Notice that like the first non-exon segment (i.e. row 1) we simulate this segment as a single site, i.e. `simDist = 1`. However, we set the recombination rate to `segLength` multiplied by the argument `recomb_rate` (i.e. 385*1E-8 = 3.85E-6).

Only three of the variables returned by `create_slimMap` are required by SLiM to simulate exon-only data: `recRate`, `mutRate`, and `endPos`. The other variables seen in the output above are used by our `read_slim` function to remap mutations to their correct positions when importing SLiM data to `R` (see section 3.2).

SLiM is written in a scripting language called Eidos. Unlike an `R` array, the first position in an Eidos array is 0. Therefore, we must shift the variable `endPos` forward 1 unit before supplying this data to SLiM.

```
# restrict output to the variables required by SLiM
slimMap <- s_map[, c("recRate", "mutRate", "endPos")]

# shift endPos up by one unit
slimMap$endPos <- slimMap$endPos - 1

# print first four rows of slimMap
head(slimMap, n = 4)
```

```
##    recRate mutRate endPos
## 1 0.00e+00   0e+00      0
## 2 1.00e-08   1e-08    354
## 3 3.85e-06   0e+00    355
## 4 1.00e-08   1e-08    464
```

We use SLiM to generate neutral mutations in exons and allowed them to accumulate over 44,000 generations, as in [4]. However, SLiM is incredibly versatile and can accommodate many different types of simulations. The creators of SLiM provide excellent resources and documentation to simulate forwards-in-time evolutionary data, which can be found at the SLiM website: https://messerlab.org/slim/.

#### 3.2 Import SLiM Data to R

To import SLiM data to `R`, we provide the `read_slim` function, which has been tested for SLiM versions 2.0-3.1. Presently, the `read_slim` function is only appropriate for data produced by the outputFull() method in SLiM. We do not support output in MS or VCF data format (i.e. produced by outputVCFsample() or outputMSSample() in SLiM).

Our `read_slim` function has four arguments:

1. `file_path`: The file path of the .txt output file created by the outputFull() method in SLiM.
2. `recomb_map`: (Optional) A recombination map of the same format as the data frame returned by `create_slimMap`. Users who followed the instructions in section 3.1, may supply the output from `create_slimMap` to this argument.
3. `keep_maf`: The largest allele frequency for retained SNVs, by default `keep_maf = 0.01`. All variants with allele frequency greater than `keep_maf` will be removed. Please note that removing common variants is recommended for large data sets due to the limitations of data allocation in R.
4. `recode_recurrent` A logical argument that indicates if recurrent SNVs at the same position should be catalogued as single observation; by default, `recode_recurrent = TRUE`.
5. `pathway_df`: (Optional) A data frame that contains the positions for each exon in a pathway of interest. This data frame must contain the variables `chrom`, `exonStart`, and `exonEnd`. *We expect that `pathwayDF` does not contain any overlapping segments. Users may combine overlapping exons into a single observation with our `combine_exons` function (see appendix for details).*

To clarify, the argument `recomb_map` is used to remap mutations to their actual locations and chromosomes. This is necessary when data have been simulated over non-contiguous regions such as exon-only data. If `recomb_map` is not provided, we assume that the SNV data have been simulated over a contiguous segment starting with the first base pair on chromosome 1.

In addition to reducing the size of the data, the argument `keep_maf` has practicable applicability. In family-based studies, common SNVs are generally filtered out prior to analysis. Users who intend to study common variants in addition to rare variants may need to run chromosome specific analyses to allow for allocation of large data sets in `R`.

When `TRUE`, the logical argument `recode_recurrent` indicates that recurrent SNVs should be recorded as a single observation. SLiM can model many types of mutations; e.g. neutral, beneficial, and deleterious. When different types of mutations occur at the same position carriers will experience different fitness effects depending on the carried mutation. However, when mutations at the same location have the same fitness effects, they represent a recurrent mutation. Even so, SLiM stores recurrent mutations separately and calculates their prevalence independently. When the argument `recode_recurrent = TRUE` we store recurrent mutations as a single observation and calculate the derived allele frequency based on their combined prevalence. This convention allows for both reduction in storage and correct estimation of the derived allele frequency of the mutation. Users who prefer to store recurrent mutations from independent lineages as unique entries should set `recode_recurrent = FALSE`.

The data frame `pathway_df` allows users to identify SNVs located within a pathway of interest when importing the data. For example, consider the apoptosis sub-pathway centered about the *TNFSF10* gene based on the 25 genes that have the highest interaction with this gene in the UCSC Genome Browser's Gene Interaction Tool [5, 6, 10]. The data for this sub-pathway are contained in the `hg_apopPath` data set.

```
# load the hg_apopPath data
data("hg_apopPath")

# View the first 4 observations of hg_apopPath
head(hg_apopPath, n = 4)
```

```
##   chrom exonStart   exonEnd
## 1     1 155689091 155689243
## 2     1 155709033 155709125
## 3     1 155709773 155709824
## 4     1 155717006 155717128
##                                                                           NCBIref
## 1 NM_001199851.1, NM_001199850.1, NM_001199849.1, NM_004632.3, NM_033657.2
## 2                                                              NM_001199849.1
## 3 NM_001199851.1, NM_001199850.1, NM_001199849.1, NM_004632.3, NM_033657.2
## 4                 NM_001199850.1, NM_001199849.1, NM_004632.3, NM_033657.2
##   gene
## 1 DAP3
## 2 DAP3
## 3 DAP3
## 4 DAP3
```

The `hg_apopPath` data set is similar the `hg_exons` data set discussed in section 3.1. However, `hg_apopPath` only catalogs the positions of exons contained in our sub-pathway.

The following code demonstrates how the `read_slim` function would be called if `create_slimMap` was used to create the recombination map for SLiM as in section 3.1.

```
# Let's suppose the output is saved in the
# current working directory and is named "slimOut.txt".
# We import the data using the read_slim function.
s_out <- read_slim(file_path  = "slimOut.txt",
                   recomb_map = create_slimMap(hg_exons),
                   pathway_df = hg_apopPath)
```

The `read_slim` function returns a list with 2 items. The first item is a sparse matrix named `Haplotypes`, which contains the haplotype data for unrelated, diploid individuals. The second item is a data frame named `Mutations`, which catalogs the SNVs contained in `Haplotypes`.

We note that importing SLiM data can be time-consuming; importing exon-only mutation data simulated over the 22 human autosomes for a sample of 10,000 individuals took approximately 4 mins on a Windows OS with an i7-4790 @ 3.60GHz and 12 GB of RAM. For the purpose of demonstration, we use the `EXhaps` and `EXmuts` and data sets provided by `SimRVSequences`.

The `EXhaps` data set represents the sparse matrix `Haplotypes` returned by `read_slim`, and the `EXmuts` data set represents the `Mutations` data frame returned by `read_slim`. We note that since these toy data sets only contain 500 SNVs they are not representative of exon-only data over the entire human genome. Instead, they include 50 SNVs that were randomly sampled from genes in the apoptosis sub-pathway, and 450 SNVs sampled from outside the pathway.

```
# import the EXmuts dataset
data(EXmuts)

# view the first 4 observations of EXmuts
head(EXmuts, n = 4)
```

```
##   colID chrom position   afreq     marker pathwaySNV
## 1     1     1  1731297 0.00875  1_1731297      FALSE
## 2     2     1  3411725 0.00005  1_3411725      FALSE
## 3     3     1  3468928 0.00050  1_3468928      FALSE
## 4     4     1 21578402 0.00025 1_21578402      FALSE
```

The `EXmuts` data frame is used to catalog the SNVs in `EXhaps`. Each row in `EXmuts` represents an SNV. The variable `colID` associates the rows in `EXmuts` to the columns of `EXhaps`, `chrom` is the chromosome the SNV resides in, `position` is the position of the SNV in base pairs, `afreq` is the derived allele frequency of the SNV, `marker` is a unique character identifier for the SNV, and `pathwaySNV` identifies SNVs located within the sub-pathway as `TRUE`.

```
# import the EXhaps dataset
data(EXhaps)

# dimensions of EXhaps
dim(EXhaps)
```

```
## [1] 20000   500
```

Looking at the output above we see that `EXhaps` contains 20,000 haplotypes with 500 SNVs.

```
# number of rows in EXmuts
nrow(EXmuts)
```

```
## [1] 500
```

Since `EXmuts` catalogs the SNVs in `EXhaps`, `EXmuts` will have 500 rows.

```
# View the first 30 mutations of the first 15 haplotypes in EXhaps
EXhaps[1:15, 1:30]
```

```
## 15 x 30 sparse Matrix of class "dgCMatrix"
##
##  [1,] . . . . . . . . . . . . . . . . . . . . . . . . . . . . . .
##  [2,] . . . . . . . . . . . . . . . . . . . 1 . . . . . . . . . .
##  [3,] . . . . . . . . . . . . . . . . . . . . . . . . . . . . . .
##  [4,] . . . . . . . . . . . . . . . . . . . . . . . . . . . . . .
##  [5,] . . . . . . . . . . . . . . . . . . . . . . . . . . . . . .
##  [6,] . . . . . . . . . . . . . . . . . . . . . . . . . . . . . .
```

```
##  [7,] . . . . . . . . . . . . . . . . . . . . . . . . . . . . . .
##  [8,] . . . . . . . . . . . . . . . . . . . . . . . . . . . . . .
##  [9,] . . . . . . . . . . . . . . . . . . . . . . . . . . . . . .
## [10,] . . . . . . . . . . . . . . 1 . . . . . . . . . . . . . . .
## [11,] . . . . . . . . . . . . . . . . . . . . . . . . . . . . . .
## [12,] . . . . . . . . . . . . . . . . . . . . . . . . . . . . . .
## [13,] . . . . . . . . . . 1 . . . . . . . . . . . . . . . . . . .
## [14,] . . . . . . . . . . . . . . . . . . . . . . . . . . . . . .
## [15,] . . . . . . . . . . . . . . . . . . . . . . . . . . . . . .
```

From the output above, we see that `EXhaps` is a sparse matrix of class dgCMatrix. The `Matrix` package [1] is used to create objects of this class. Entries of `1` represent a mutated allele, while the wild type is represented as '`.`'. Recall that we retain SNVs with derived allele frequency less than or equal to `keep_maf`; hence, the majority of entries represent the wild type allele.

#### 3.3 Select Pool of Causal Variants

Our next task is to decide which mutations will be modeled as causal variants. We will focus on a pathway approach, which allows for causal variation in a pathway, or a set of related genes. This approach may also be used to model families which segregate different rare variants in the same gene. Furthermore, we note that implementation of a single causal variant is equivalent to a pathway containing a single gene with a single mutation.

For `SimRVpedigree`'s ascertainment process to be valid we require that the probability that a random individual carries a risk allele is small, i.e $\leq 0.002$ [9]. Consider a set of $n$ causal SNVs with derived allele frequencies $p_1, p_2, ..., p_n$, and let $p_{tot} = \sum_{i=1}^{n} p_i$. For sufficiently rare SNVs, we may compute the cumulative carrier probability as $p_{tot}^2 + 2 * p_{tot}(1 - p_{tot})$. To satisfy the condition $p_{carrier} \leq 0.002$, we require that $1 - (1 - p_{tot})^2 \leq 0.002$, or that $p_{tot} \leq 0.001$.

We now demonstrate how users may create a new variable that identifies causal SNVs.

```
# Recall that the variable pathwaySNV, which was created in
# section 3.2, is TRUE for any SNVs in the pathway of interest.
# Here we tabulate the derived allele frequencies
# of the SNVs located in our pathway.
table(EXmuts$afreq[EXmuts$pathwaySNV == TRUE])
```

```
##
##    5e-05    1e-04 0.00015    2e-04 0.00035    4e-04 0.00045    5e-04 0.00055
##       12        4       6        1       1        1       1        1       1
## 0.00065    7e-04 0.00075    8e-04    9e-04 0.00095    0.001 0.00125   0.0014
##        1        1       1        1        2       1        1       1        1
## 0.00145 0.00165   0.002 0.00205   0.0021 0.00365 0.00505 0.00515   0.0052
##       1       1       1       1        1       1       1       1        1
## 0.00585 0.00635 0.00925
##       1       1       1
```

The output above tabulates variants in our pathway by their derived allele frequencies. For example, our pathway contains 12 SNVs with derived allele frequency 5e-05, 4 SNVs with derived allele frequency 1e-04, 6 SNVs with derived allele frequency 0.00015, and so on.

To obtain a pool of variants with $p_{tot} \sim 0.001$, let's choose the 12 SNVs with derived allele frequency 5e-05, and 4 SNVs with derived allele frequency 1e-4 to be our causal SNVs, so that $p_{tot} = 12(5 \times 10^{-5}) + 4(1 \times 10^{-4}) = 0.001$.

```
# Create the variable 'is_CRV', which is TRUE for SNVs
# in our pathway with derived allele frequency 5e-05 or 1e-04,
# and FALSE otherwise
```

```r
EXmuts$is_CRV <- EXmuts$pathwaySNV & EXmuts$afreq %in% c(5e-5, 1e-04)

# verify that sum of the derived allele
# frequencies of causal SNVs is 0.001
sum(EXmuts$afreq[EXmuts$is_CRV])
```

```
## [1] 0.001
```

```r
# determine the number of variants in our pool of causal variants
sum(EXmuts$is_CRV)
```

```
## [1] 16
```

From the output above we see that we have selected a pool of 16 causal variants from our pathway with a cumulative allele frequency of 0.001.

```r
# view first 4 observations of EXmuts
head(EXmuts, n = 4)
```

```
##   colID chrom position   afreq     marker pathwaySNV is_CRV
## 1     1     1  1731297 0.00875  1_1731297      FALSE  FALSE
## 2     2     1  3411725 0.00005  1_3411725      FALSE  FALSE
## 3     3     1  3468928 0.00050  1_3468928      FALSE  FALSE
## 4     4     1 21578402 0.00025 1_21578402      FALSE  FALSE
```

Observe from the output above that `EXmuts` now contains the variable `is_CRV`, which identifies potential causal rare variants. The `sim_RVstudy` function (discussed in section 5) samples a single causal variant for each family, according to its allele frequency in the population, from the SNVs for which `is_CRV = TRUE`. Hence, different families may segregate different causal rare variants. Upon identifying the familial cRV we then sample haplotypes for each founder from the distribution of haplotypes conditioned on the founder's cRV status. This ensures that the cRV is introduced by the correct founder.

### 4 Prepare Pedigree Data

The R package `SimRVPedigree` [8] is used to simulate pedigrees ascertained for multiple disease-affected relatives. The following example demonstrates how users could use `SimRVPedigree` to simulate five pedigrees ascertained for three or more disease-affected relatives in parallel with the `doParallel`[7] and `doRNG`[2] packages.

```r
# load the SimRVPedigree library
library(SimRVPedigree)

# Create hazard object from AgeSpecific_Hazards data
data(AgeSpecific_Hazards)
my_HR = hazard(AgeSpecific_Hazards)

# load libraries needed to simulate pedigrees in parallel.
library(doParallel)
library(doRNG)

npeds <- 5    #set the number of pedigrees to generate

cl <- makeCluster(2)   # create cluster
registerDoParallel(cl) # register cluster
```

```r
# simulate a sample of five pedigrees using foreach
study_peds = foreach(i = seq(npeds), .combine = rbind,
                  .packages = c("SimRVPedigree"),
                  .options.RNG = 844090518
) %dorng% {
  # Simulate pedigrees ascertained for at least three disease-affected individuals,
  # according to the age-specific hazard rates in the `AgeSpecific_Hazards` data
  # set, ascertained from 1980 to 2010, with start year spanning
  # from 1900 to 1920, stop year set to 2018, and with genetic relative-risk 50.
  sim_RVped(hazard_rates = my_HR,
            GRR = 50, FamID = i,
            RVfounder = TRUE,
            founder_byears = c(1900, 1920),
            ascertain_span = c(1980, 2010),
            stop_year = 2018,
            recall_probs = c(1, 0.5, 0),
            num_affected = 3)[[2]]}

stopCluster(cl) #shut down cluster
```

For the purpose of demonstration, we will use the `study_peds` data set provided by `SimRVSequences`.

```r
# import study peds
data("study_peds")

# Plot the pedigree with FamID 3
plot(study_peds[study_peds$FamID == 3, ],
     ref_year = 2018)
```

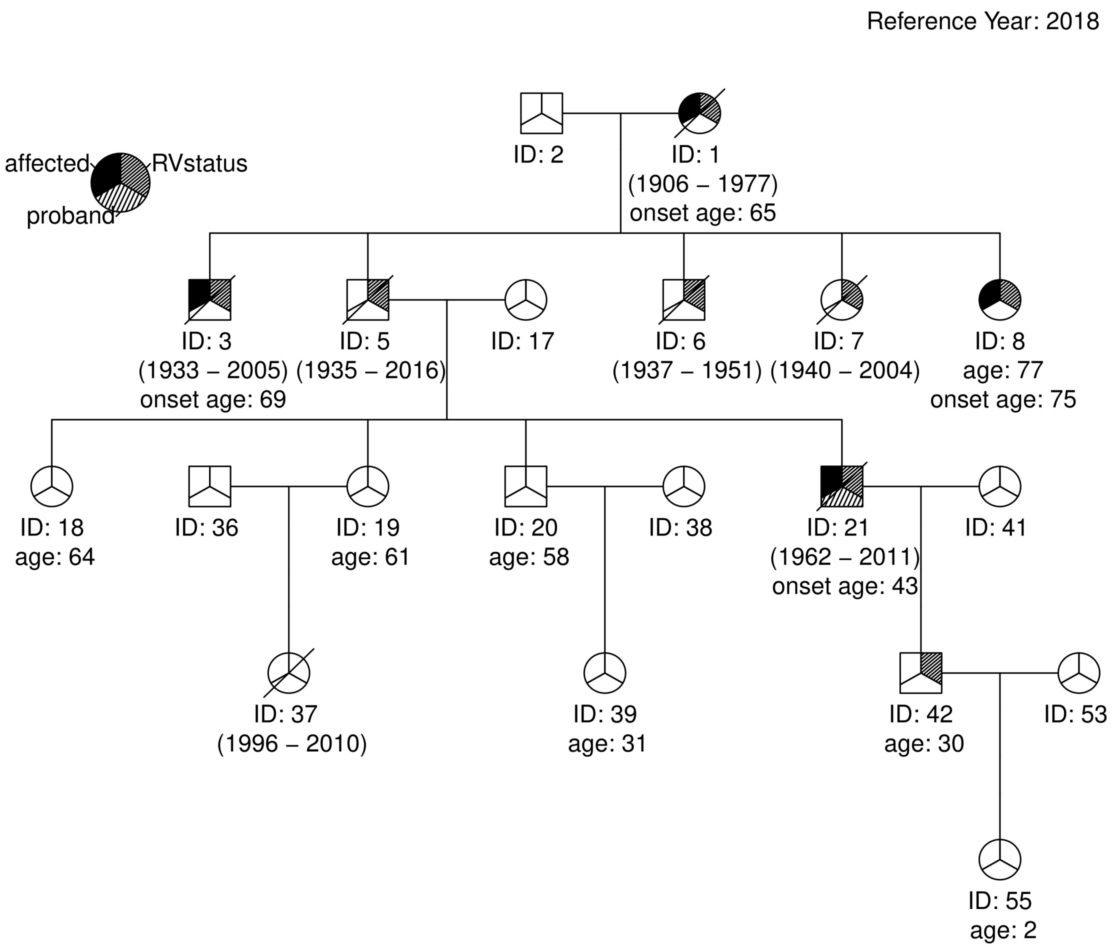

According to the legend, disease-affected individuals (IDs 1, 3, 8, and 21) are indicated by a solid shading in the upper-left third of their symbol. The proband (ID 21) is indicated by shading in the lower portion of his symbol. Individuals who have inherited the cRV (IDs 1, 3, 5, 6, 7, 8, 21 and 42) are indicated by shading in the upper-right portion of their symbol.

The seed founder (ID 1) and all of his descendants have age data relative to the reference year, which is displayed below their symbols. If a descendant has died by the reference year (IDs 1, 3, 5, 6, etc.) their year of birth and death are displayed in parentheses, and their symbol will have a diagonal line over it. Descendants who are still alive at the reference year have an age identifier. Descendants who have experienced disease-onset by the reference year will have a disease-onset age identifier.

```
# View the first 4 rows of study_peds
head(study_peds, n = 4)
```

```
##   FamID ID sex dadID momID affected DA1 DA2 birthYr onsetYr deathYr RR
## 1     1  1   1    NA    NA     TRUE   1   0    1902    1985    1987 50
## 2     1  2   0    NA    NA    FALSE   0   0      NA      NA      NA  1
## 3     1  3   0     2     1    FALSE   0   1    1922      NA    1942 50
```

```
## 4     1  4   1     2     1     TRUE   0   1    1925    1991    1991 50
##   available Gen proband
## 1      TRUE   1   FALSE
## 2     FALSE   1   FALSE
## 3      TRUE   2   FALSE
## 4      TRUE   2    TRUE
```

The columns displayed in the output above are described as follows:

1. `FamID`: family identification number
2. `ID`: individual identification number
3. `sex`: sex identification variable: `sex = 0` for males, and `sex = 1` females.
4. `dadID`: identification number of father
5. `momID`: identification number of mother
6. `affected`: disease-affection status: `affected = TRUE` if individual has developed disease, and `FALSE` otherwise.
7. `DA1`: paternally inherited allele at the cRV locus: `DA1 = 1` if the cRV is inherited, and `0` otherwise.
8. `DA2`: maternally inherited allele at the cRV locus: `DA2 = 1` if the cRV is inherited, and `0` otherwise.
9. `birthYr`: the individual's birth year.
10. `onsetYr`: the individual's year of disease onset, when applicable, and `NA` otherwise.
11. `deathYr`: the individual's year of death, when applicable, and `NA` otherwise.
12. `RR`: the individual's relative-risk of disease.
13. `available`: availability status: `available = TRUE` if individual is recalled by the proband, and `FALSE` if not recalled or a marry-in.
14. `Gen`: the individual's generation number relative to the eldest pedigree founder. That is, the seed founder will have `Gen = 1`, his or her offspring will have `Gen = 2`, etc.
15. `proband`: a proband identifier: `proband = TRUE` if the individual is the proband, and `FALSE` otherwise.

Not all of the variables above are required to simulate SNV data; the required variables are: `FamID`, `ID`, `sex`, `dadID`, `momID`, `affected`, `DA1`, and `DA2`. Please note, if `DA1` and `DA2` are not provided, the pedigree is assumed to be fully-sporadic, i.e. `DA1 = DA2 = 0` for all members.

To learn more about simulating pedigrees with `SimRVPedigree` please refer to `SimRVPedigree`'s documentation and vignette.

### 5 Simulate SNV data for Pedigrees

Simulation of sequence data for ascertained pedigrees is accomplished with the `sim_RVstudy` function. We will illustrate the usage of this function in section 5.1. The `sim_RVstudy` function returns an object of class `famStudy`. In section 5.2 we discuss objects class `famStudy` in detail. Finally, in section 5.3 we demonstrate the usage and output of the `summary` function for objects of class `famStudy`.

#### 5.1 The `sim_RVstudy` function

Simulating SNV data for a sample of ascertained pedigrees is achieved with our `sim_RVstudy` function. This function has the following required and optional arguments:

Required Arguments:

1. `ped_files` A data frame of pedigrees. This data frame must contain the variables: `FamID`, `ID`, `sex`, `dadID`, `momID`, `affected`, `DA1`, and `DA2`. If `DA1` and `DA2` are not provided we assume the pedigrees are fully sporadic. See section 4 for additional details regarding these variables.
2. `haplos` A sparse matrix of haplotype data, which contains the haplotypes for unrelated individuals representing the founder population. Rows are assumed to be haplotypes, while columns represent

SNVs. If `read_slim` was used to import data, users may supply the sparse matrix `Haplotypes` returned by `read_slim`.

3. `SNV_map` The rows of the data frame `SNV_map` provide information on the SNVs in `haplos.` If `read_slim` was used to import data, the data frame `Mutations` returned by this function is of the proper format for `SNV_map` (see section 3.2). We note that users must add the variable `is_CRV` to this data frame, as discussed in section 3.3.

Optional Arguments:

4. `affected_only` A logical argument. When `affected_only = TRUE`, we only simulate SNV data for the affected individuals and the family members that connect them along a line of descent. When `affected_only = FALSE`, SNV data is simulated for the entire pedigree. By default, `affected_only = TRUE`.
5. `remove_wild` A logical argument that determines if SNVs not carried by any member of the study should be removed from the data. By default, `remove_wild = TRUE`.
6. `pos_in_bp` A logical argument which indicates if the positions in `SNV_map` are listed in base pairs. By default, `pos_in_bp = TRUE`. If the positions in `SNV_map` are listed in centiMorgans set `pos_in_bp = FALSE` instead.
7. `gamma_params` The respective shape and rate parameters of the gamma distribution used to simulate the distance between chiasmata. By default, `gamma_params = c(2.63, 2*2.63)`, as in [11].
8. `burn_in` The “burn-in” distance in centiMorgans, as defined by [11], which is required before simulating the location of the first chiasmata with interference. By default, `burn_in = 1000`.

Assuming readers have followed the steps outlined in sections 3 and 4, we may execute the `sim_RVstudy` function as follows.

```
set.seed(11956)
# simulate SNV data using sim_RVstudy
study_seq <- sim_RVstudy(ped_files = study_peds,
                         SNV_map = EXmuts,
                         haplos = EXhaps)

# determine the class of study seq
class(study_seq)
```

```
## [1] "famStudy" "list"
```

From the output above, we see that the `sim_RVstudy` function returns an object of class `famStudy`, which inherits from the `list` class. More specifically, a `famStudy` object is a list that contains the following four items:

- a data frame named `ped_files`,
- a sparse matrix named `ped_haplos`,
- a data frame named `haplo_map`,
- and a data frame named `SNV_map`.

We will discuss each of the four items contained in the `famStudy` object in the following section.

#### 5.2 Objects of class `famStudy`

##### 5.2.1 The `ped_files` data frame

The data frame `ped_files` contained in the `famStudy` object is a potentially reduced set of pedigree files for the original families. Recall that, by default `affected_only = TRUE`, so the pedigrees supplied to `sim_RVstudy` are reduced to contain only the affected individuals and the individuals who connect them along a line of descent.

```r
# plot the pedigree returned by sim_RVseq for family 3.
# Since we used the affected_only option the pedigree has
# been reduced to contain only disease-affected relatives.
plot(study_seq$ped_files[study_seq$ped_files$FamID == 3, ],
     location = "bottomleft")
```

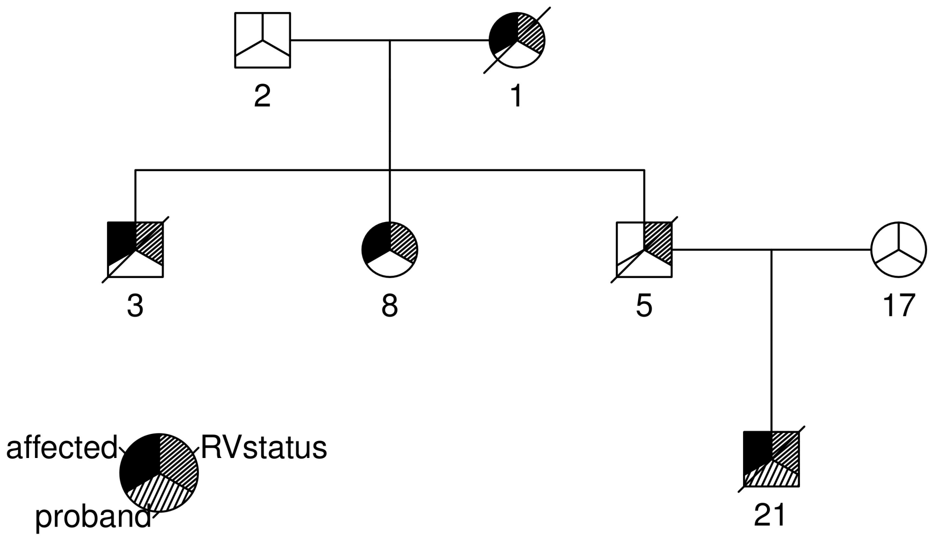

Comparing the plot above to the plot of the same family in section 4, we can see that the number of relatives for whom we simulate SNV data is far less than the total number of relatives in the ascertained pedigree.

###### 5.2.2 The `ped_haplos` matrix

Next, we look at the sparse matrix, `ped_haplos`, which is contained in the `famStudy` object. This sparse matrix contains simulated haplotype data.

```r
# View the first 15 haplotypes in ped_haplos
study_seq$ped_haplos[1:15, ]
```

```
## 15 x 25 sparse Matrix of class "dgCMatrix"
##
##  [1,] . . . 1 . . . . . . . . . . . . . . . . . . . . .
##  [2,] . . . . . . . . . . . . . . . . . . . . . . . . .
##  [3,] . . . . . . . . . . . . . . . . 1 . . . . . . . .
##  [4,] 1 . . . 1 . . . . . . . . . . . . . . . . . . . .
##  [5,] . . . . . . . . . . . . . . . . . . . . . . . . .
##  [6,] . . . 1 . . . . . . . . . . . . . . . . . . . . .
##  [7,] 1 . . . 1 . . . . . . . . . . . 1 . . . . . . . .
```

```
##  [8,] . . . 1 . . . . . . . . . . . . . . . . . . . . .
##  [9,] . . . . . . . . . . . . . . . . . . . . . . . . .
## [10,] . . . 1 . . . . . . . . . . . . . . . . . . . . .
## [11,] . 1 . . . . . . . . . . . . . . . . . . . 1 . . .
## [12,] . . . . . . . . . . . . . . . . . . . . . . . . .
## [13,] . . . . . . . . . . . . . . . . . . . . . . . . .
## [14,] . . . . . . . . . . . . . . . . . . 1 . . . . . .
## [15,] . . . . . . . . . . . . . . . . . . 1 . . . . . .
```

The `ped_haplos` matrix contains the haplotype data for each individual in the `ped_files` output.

```
# Determine the dimensions of ped_haplos
dim(study_seq$ped_haplos)
```

```
## [1] 62 25
```

```
# Determine the dimensions of EXhaps
dim(EXhaps)
```

```
## [1] 20000   500
```

Notice that `ped_haplos` only has 24 columns (i.e. mutations) even though `EXhaps`, which was supplied to `sim_RVstudy`, has a total of 500 columns. This occurs when the `remove_wild` setting is used to remove superfluous SNVs from our data. That is, when no one in the study carries a mutated allele at an SNV locus, that marker is removed from the data. Since we are focused on rare variation, and since `EXhaps` and `study_peds` are small toy data sets the reduction in data is quite significant.

##### 5.2.3 The SNV_map data frame

In this subsection we discuss the `SNV_map` data frame contained in the `famStudy` object. Recall that one of `sim_RVstudy`'s arguments is also named `SNV_map`. When the `remove_wild` argument of `sim_RVstudy` is set to `FALSE`, the data frame supplied to `SNV_map` will be returned unchanged. However, when `remove_wild` is set to `TRUE`, the `SNV_map` contained in the `famStudy` object will be a reduced data frame that contains only SNVs that are carried by at least one study participant.

```
# View the first 4 observations in SNV_map
head(study_seq$SNV_map, n = 4)
```

```
##   colID chrom  position   afreq      marker pathwaySNV is_CRV
## 1     1     1  45043357 0.00310  1_45043357      FALSE  FALSE
## 2     2     1 236599491 0.00895 1_236599491      FALSE  FALSE
## 3     3     2   3960458 0.00535   2_3960458      FALSE  FALSE
## 4     4     2 201284953 0.00010 2_201284953       TRUE   TRUE
```

The output `SNV_map` catalogs the SNVs, i.e. columns, in `ped_haplos`. For example, the first column in `ped_haplos` is a SNV located on chromosome 1 at position 45043357.

##### 5.2.4 The haplo_map data frame

Next, we look at the `haplo_map` data frame contained in the `famStudy` object. The `haplo_map` data frame is used to map the haplotypes (i.e. rows) in `ped_haplos` to the individuals in `ped_files`.

```
# view the first observations in haplo_map
head(study_seq$haplo_map)
```

```
##   FamID ID affected      FamCRV
## 1     1  1     TRUE 2_201284953
```

```
## 2     1  1     TRUE 2_201284953
## 3     1  2    FALSE 2_201284953
## 4     1  2    FALSE 2_201284953
## 5     1  4     TRUE 2_201284953
## 6     1  4     TRUE 2_201284953
```

Notice that in the output above individual 1 from family 1 is listed in rows 1 and 2, and individual 2 from family 1 is listed in rows 3 and 4. This is because the genetic material inherited from each parent is stored in its own row. **By convention, the first row represents the paternally inherited haplotype, while the second row represents the maternally inherited haplotype**. Additionally, note the variable `FamCRV` contained in `haplo_map`, which identifies each family's causal rare variant by marker name.

The following code demonstrates how to reduce `haplo_map` to quickly identify the cRV for each family.

```
# to quickly view the causal rare variant for
# each family we supply the appropriate columns of
# haplo_map to unique
unique(study_seq$haplo_map[, c("FamID", "FamCRV")])
```

```
##    FamID      FamCRV
## 1      1 2_201284953
## 11     2      no_CRV
## 19     3  8_23081797
## 33     4 2_201284953
## 41     5 10_89008915
```

From the output above, we see that families 1 and 4 segregate variant `2_201284953`, family 2 does not segregate any SNV in the pool of causal rare variants, family 3 segregates variant `10_89016102`, and family 5 segregates variant `10_89015798`. Note that family 2 does not segregate any of the SNVs in our pool of causal rare variants because it is a fully sporadic pedigree; i.e. there is no genetic predisposition to disease in this pedigree.

##### 5.3 The `summary.famStudy` function

To obtain summary information for the simulated sequence data we supply objects of class `famStudy` to the `summary` function. Recall that the output of the `sim_RVstudy` function is an object of class `famStudy`; hence, users may supply the output of `sim_RVstudy` directly to `summary`.

```
# supply the famStudy object, returned by
# sim_RVstudy, to the summary function
study_summary <- summary(study_seq)

# determine the class of study_summary
class(study_summary)
```

```
## [1] "list"
```

```
# view the names of the items in study_summary
names(study_summary)
```

```
## [1] "fam_allele_count" "pathway_count"
```

From the output above, we see that when supplied an object of class `famStudy` the `summary` function returns a list containing two items: `fam_allele_count` and `pathway_count`. The item `fam_allele_count` is a matrix that includes counts of the SNVs shared among the disease-affected relatives in each pedigree.

```
# view fam_allele_count
study_summary$fam_allele_count
```

```
##      FamID 1_45043357 1_236599491 2_3960458 2_201284953 3_47610321
## [1,]     1          1           0         0           4          1
## [2,]     2          0           3         0           0          0
## [3,]     3          0           0         3           0          0
## [4,]     4          0           0         0           3          0
## [5,]     5          0           0         0           0          0
##      3_49154162 3_129583578 4_2249748 4_168989553 5_77691934 5_115614755
## [1,]          0           0         0           0          0           0
## [2,]          0           0         0           0          0           0
## [3,]          1           1         0           1          0           0
## [4,]          0           0         0           0          0           1
## [5,]          0           0         1           0          1           0
##      7_101314288 8_23081797 8_144514988 10_89008915 12_8946339 12_26936616
## [1,]           0          0           0           0          1           0
## [2,]           0          0           0           0          0           0
## [3,]           2          4           2           0          0           2
## [4,]           0          0           0           0          0           0
## [5,]           0          0           0           4          0           0
##      15_63129900 17_58007302 21_33437240 21_33449698 22_50460703
## [1,]           0           0           0           0           0
## [2,]           1           2           0           0           0
## [3,]           0           0           0           3           2
## [4,]           0           0           2           0           0
## [5,]           0           0           0           0           0
```

Looking at the output above, we see that only one affected relative in family 1 carries a copy of the first SNV, 1_45043357. It is noteworthy that the disease-affected relative in families 1 and 4 carry so many copies of SNV 2_201284953; not surprisingly, this is the causal rare variant for these two families.

The item pathway_count is a data frame that catalogs the total number of SNVs observed in disease-affected study participants.

```
# view pathway_count
study_summary$pathway_count
```

```
##    chrom  position      marker total is_CRV pathwaySNV
## 1      1  45043357  1_45043357     1  FALSE      FALSE
## 2      1 236599491 1_236599491     3  FALSE      FALSE
## 3      2   3960458   2_3960458     3  FALSE      FALSE
## 4      2 201284953 2_201284953     7   TRUE       TRUE
## 5      3  47610321  3_47610321     1  FALSE      FALSE
## 6      3  49154162  3_49154162     1  FALSE      FALSE
## 7      3 129583578 3_129583578     1  FALSE      FALSE
## 8      4   2249748   4_2249748     1  FALSE      FALSE
## 9      4 168989553 4_168989553     1  FALSE      FALSE
## 10     5  77691934  5_77691934     1  FALSE      FALSE
## 11     5 115614755 5_115614755     1  FALSE      FALSE
## 12     7 101314288 7_101314288     2  FALSE      FALSE
## 14     8  23081797  8_23081797     4   TRUE       TRUE
## 15     8 144514988 8_144514988     2  FALSE      FALSE
## 16    10  89008915 10_89008915     4   TRUE       TRUE
## 17    12   8946339  12_8946339     1  FALSE      FALSE
## 18    12  26936616 12_26936616     2  FALSE      FALSE
## 19    15  63129900 15_63129900     1  FALSE      FALSE
## 22    17  58007302 17_58007302     2  FALSE      FALSE
## 23    21  33437240 21_33437240     2  FALSE       TRUE
```

```
## 24    21   33449698 21_33449698      3  FALSE        TRUE
## 25    22   50460703 22_50460703      2  FALSE       FALSE
```

From `pathway_count`, we see that there are several SNVs carried by more than 1 disease-affected study participants; however, only 5 of the SNVs carried by more than one individual reside in the pathway of interest.

### 6 SimRVSequences Model

In this section we describe the model assumptions employed by `SimRVSequences` to simulate single-nucleotide variant (SNV) data for pedigrees.

1. Given a sample of ascertained pedigrees we allow families to segregate different rare variants, but assume that within a family genetic cases are due to a single, **causal rare variant (cRV)** that increases disease susceptibility. We assume that only one copy of the familial cRV is introduced to the pedigree and that the cRV status of every pedigree member is known. The `R` package `SimRVPedigree` [8] may be used to simulate pedigrees that meet these criteria (see section 4).

2. Recall that users are expected to specify a pool of cRVs which may represent cRVs located in a pathway, or different cRVs in the same gene. The size of the pool is entirely up to the user: it can contain many SNVs or only a single SNV. However, we urge those who are using `SimRVPedigree` in conjunction with `SimRVSequences` to choose cRVs so that their cumulative carrier probability is less than or equal to 0.002 [9]. From this pool we sample a single cRV for each pedigree with replacement.

3. Founder haplotypes are sampled from a user-specified, population distribution of haplotypes, conditional on their cRV status.
    - For the founder who introduces the cRV we sample one haplotype from the haplotypes that carry the familial cRV and the other from the haplotypes that do not carry ANY of the cRVs in our pool. This ensures that this founder introduces only a single copy of the familial cRV.
    - For the pedigree founders who do NOT introduce a cRV we sample their haplotypes from the haplotypes that do not carry ANY of the cRVs in our pool. Hence, fully sporadic families will not segregate any cRVs, nor will any non-carrier individual who marries into a genetic family.

4. We use a conditional gene-drop algorithm to model inheritance from parent to offspring, which can be described as follows.

- First, we simulate the formation of a parent's gametes.
    1. Following [11], we model genetic recombination with chiasmata interference. To accomplish this the distance between chiasmata is modelled by a gamma distribution. Additionally, we use a "burn-in" process before the start of the chromosome to simulate the location of the first chiasmata. The default parameter settings for the gamma distribution are `shape = 2.63` and `rate = 2*2.63`, and `burn_in = 1000`. To model recombination without chiasmata interference, i.e. according to Haldane's model, set `shape = 1` and `rate = 2` and `burn_in = 0`. **Please note, presently `SimRVSequences` only permits simulation of recombination in autosomes, i.e. non-sex chromosomes.**
    2. We assume no chromatid interference so that non-sister chromatids are equally likely to participate in a crossover event.
    3. To simulate the formation of gametes we assume that homologous chromatids are assigned to one of four gamete cells with equal probability. This assignment occurs independently for non-homologous chromosomes.

- We use a conditional gene drop algorithm to determine which of the four gametes is transmitted from parent to offspring by considering the cRV status of the parent and the offspring, as follows.

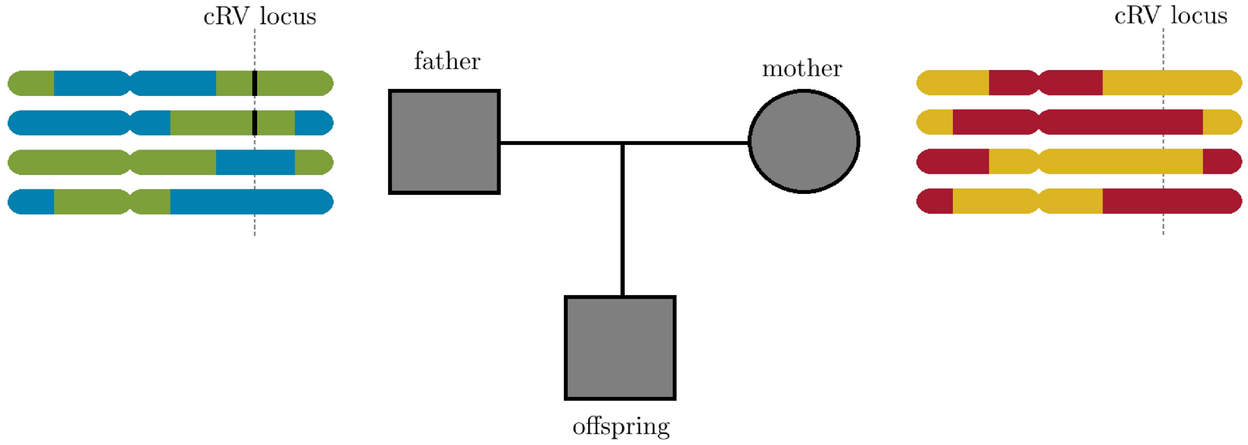

Figure 2: **Figure 2.** A family trio with possible maternal and paternal recombinant chromatids for a diploid organism with a single set of homologous chromosomes. The cRV locus is indicated by a dashed vertical line over the chromatids, and the cRV is indicated by a black vertical bar within the chromatids. The top two chromatids of the father contain the only copies of the cRV.

- Case 1: If **both the parent and offspring carry the cRV** we sample the inherited gamete from the two possible parental gametes that carry the cRV. For example, in the father to offspring transmission in Figure 2, we choose from the paternal gametes represented by the top two chromatids.
- Case 2: If the **parent carries the cRV but the offspring does not**, we sample the inherited gamete from the two possible parental gametes that do not carry the cRV. For example, in the father to offspring transmission in Figure 2, we choose from the paternal gametes represented by the bottom two chromatids.
- Case 3: If the **parent is not a carrier of the cRV**, then the cRV status of the offspring is irrelevant for this parent's transmission because the offspring cannot inherit it from this parent. In this scenario, we sample the inherited gamete from the four parental gametes with equal probability. For example, this would be the case in the mother to offspring transmission in figure 2, since she is not a carrier of the cRV.

We use these assumptions in our algorithm to simulate sequence data for pedigrees (see introduction.)

### 7 Method Overview

The methods provided by `SimRVSequences` are described as follows.

- Methods to assist with creating the recombination map for SLiM:
  1. `combine_exons`: This function will be useful to users who would like to combine overlapping exon observations into a single observation. Upon supplying a data frame which contains overlapping exon segments, this function will return a data frame with combined segments (see appendix).

  2. `create_slimMap`: This function is used both before and after simulating SLiM data. Prior to simulating the SNV data with SLiM, this function can be used to create the recombination map required by SLiM to simulate exon-only SNV data (see section 3.1). Additionally, if this function was used to create the recombination map users may then supply the output of this function to `read_slim` to remap mutations to their correct locations when importing SLiM data to `R` (see section 3.2).
- Method to assist with importing SLiM data to `R`.

3. `read_slim`: This function is used to import to `R` data stored to a .txt file by SLiM's outputfull() method (see section 3.2).

- Methods to assist with simulation and basic manipulation of SNV data for pedigrees
  4. `sim_RVstudy`: This function is used to simulate the SNV data for a sample of pedigrees (see section 5.1).
  5. `summary.famStudy`: This function is used to obtain a summary of the SNVs shared by the disease-affected relatives in each pedigree (see section 5.3).

### 8 Timing

**All of the following timing estimates were performed on a Windows OS with an i7-4790 @ 3.60GHz and 12 GB of RAM.**

Importing genome-wide, exon-only SNV data for 10,000 diploid individuals from the .txt file produced by SLiM's outputfull() method, using the `read_slim` function required approximately 4 minutes. The imported data originally contained a total of 306,825 SNVs over 20,000 haplotypes.

For timing of `sim_RVstudy`, we explore the effects of the `affected_only` and `remove_wild` arguments. When `affected_only = TRUE`, sequence data is only simulated for disease-affected relatives and the family members that connect them along a line of descent, otherwise if `affected_only = FALSE` sequence data is simulated for the entire pedigree. When `remove_wild = TRUE` the size of the data is reduced by removing from the data SNVs not carried by any study participant; otherwise if `remove_wild = FALSE` no data reduction occurs.

The following timings were performed on a sample of 100 pedigrees, each of which contained at least 3 relatives affected by a lymphoid cancer. To simulate these pedigrees we used the same settings as in the parallel processing example of section 4. For timing, we used genome-wide, exon-only SNV data for 10,000 diploid individuals simulated by SLiM. After reducing the SLiM data to contain SNVs with derived allele frequency less than or equal to 0.01, there were a total of 178,997 SNVs in our data. We chose a cRV pool of size 20 at random from the SNVs with a derived allele frequency of 5e-05 located in the sub-apoptosis pathway described by the `hg_apopPath` data set (see section 3.2 and 3.3).

| Number of Pedigrees | `affected_only` | `remove_wild` | time (in minutes) |
|---|---|---|---|
| 100 | TRUE | TRUE | 0.976 |
| 100 | TRUE | FALSE | 1.267 |
| 100 | FALSE | TRUE | 2.032 |
| 100 | FALSE | FALSE | 3.051 |

For the sample of 100 pedigrees used to time `sim_RVstudy`, setting `affected_only = FALSE` resulted in simulation of sequence data for a total of 1473 individuals, when `affected_only = TRUE` the data were reduced to contain a total of 487 individuals. On average, this resulted in a reduction of 9.86 individuals from each pedigree.

### 10 Appendix

The `combine_exons` function is used to combine overlapping exons into single segments.

This function has only one argument called `exon_data`, which is a data frame that includes the following variables

1. `chrom` a chromosome identifier,
2. `exonStart` the first position of the exon in base pairs, and
3. `exonEnd` the last position of the exon in base pairs.

Usage of the `combine_exons` function is illustrated below.

```
# create an example data frame that contains the
# the variables: chrom, exonStart, and exonEnd
exDat <- data.frame(chrom     = c(1, 1, 1, 2, 2, 2),
                    exonStart = c(1, 2, 5, 1, 3, 3),
                    exonEnd   = c(3, 4, 7, 4, 5, 6))

# View exDat data set
exDat
```

```
##   chrom exonStart exonEnd
## 1     1         1       3
## 2     1         2       4
## 3     1         5       7
## 4     2         1       4
## 5     2         3       5
## 6     2         3       6
```

From the output above, we see that the first two exons in chromosome one overlap (i.e. the exons with ranges [1, 3] and [2, 4]), as do all three exons in chromosome two.

```
# supply exDat to combine_exons
# and view the results
combine_exons(exDat)
```

```
##   chrom exonStart exonEnd
## 1     1         1       7
## 2     2         1       6
```

After supplying the data to `combine_exons` we see that the segments [1, 3] and [2, 4] on chromosome one have been combined in to a single segment: [1, 4]. Similarly, the three overlapping exons from chromosome two: [1, 4], [3, 5], and [3, 6], have now been combined into the segment [1, 6].
