## Supplementary Table for "SimRVSequences: an R package to simulate genetic sequence data for pedigrees"

### Feature Comparison for Sequence Data Simulation in Pedigrees

| | `SimRVSequences` | `sim1000G` | RarePedSim |
|---|---|---|---|
| programming language | R | R | Python and C++ |
| exon-only option | YES | NO | NO |
| total number of variants | tested up to 250,000[1] | tested up to 25,000[2] | gene/region level |
| simulate chromosomes simultaneously | YES | NO | NO |
| recombination with chiasmata interference | YES | YES | NO |
| simulate founder genotypes | NO[3] | YES | YES |
| conditional gene-drop[4] | YES | NO | YES[5] |
| simulate genotype given phenotype | NO | NO | YES |
| pathway implementation | YES | NO | NO |
| genetic heterogeneity within pedigree | NO | NO | YES |
| genetic heterogeneity of disease | YES | NO | YES |
| accepts standard pedigree format, e.g. linkage file | YES | NO | YES |
| flexible variant types | NO[6] | YES | YES |

[1]Windows OS, i7-4790 @ 3.60GHz and 12GB of RAM.

[2]Windows OS, i7-4790 @ 3.60GHz and 12GB of RAM.

[3]See section 3.4 of main text for details.

[4]However, we provide methods to assist with simulating founder genotypes with SLiM.

[5]User must set phenotypes to observed cRV status and penetrance to 1.

[6]Users are limited to simulation of single nucleotide variants (SNVs).
